# Single cell BCR and transcriptome analysis after respiratory virus infection reveals spatiotemporal dynamics of antigen-specific B cell responses

**DOI:** 10.1101/2020.08.24.264069

**Authors:** Nimitha R. Mathew, Jayalal K. Jayanthan, Ilya Smirnov, Jonathan L. Robinson, Hannes Axelsson, Sravya S. Nakka, Aikaterini Emmanouilidi, Paulo Czarnewski, William T. Yewdell, Cristina Lebrero-Fernández, Valentina Bernasconi, Ali M. Harandi, Nils Lycke, Nicholas Borcherding, Jonathan W. Yewdell, Victor Greiff, Mats Bemark, Davide Angeletti

## Abstract

B cell responses are a critical component of anti-viral immunity. However, a comprehensive picture of antigen-specific B cell responses, differentiation, clonal proliferation and dynamics in different organs after infection is lacking. Here, we combined single-cell RNA sequencing with single-cell B cell receptor (BCR) characterization of antigen-specific cells in the draining lymph nodes, spleen and lungs after influenza infection. We identify several novel B cell subpopulations forming after infection and find organ-specific differences that persist over the course of the response. We discover important transcriptional differences between memory cells in lungs and lymphoid organs and describe organ-restricted clonal expansion. Strikingly, by combining BCR mutational analysis, monoclonal antibody expression and affinity measurements we find no differences between germinal center (GC)-derived memory and plasmacells, at odds with an affinity-based selection model. By linking antigen-recognition with transcriptional programming, clonal-proliferation and differentiation, these finding provide important advances in our understanding of antiviral B cell immunity.

## INTRODUCTION

Viral respiratory infections caused by influenza-, orthopneumo- or corona-virus are major concerns worldwide. B cell derived antibodies (Abs) are a central feature of adaptive immunity to viruses. Abs can greatly reduce viral pathogenicity in primary infections and can provide complete protection against disease causing reinfections (Lam and Baumgarth, 2019). Influenza A virus (IAV) is a highly prevalent respiratory virus that causes significant morbidity and mortality in humans (Iuliano et al., 2017). Intranasal (i.n.) infection with IAV initiates B cell responses in several organs characterized by a robust early extrafollicular plasmablast (PB) response, followed by persistent germinal center (GC) formation in the draining mediastinal lymph nodes (mln) and diffuse memory B cell (Bmem) dispersion across several organs (Angeletti et al., 2017; Boyden et al., 2012; Frank et al., 2015; Joo et al., 2008; Rothaeusler and Baumgarth, 2010). Respiratory virus infection can also promote circulating blood cells to generate inducible bronchus-associated lymphoid tissues (iBALTs) in the lung parenchyma (Moyron-Quiroz et al., 2004) exemplified by the formation of GC-like structures in mouse lungs by 14 days post infection (dpi) with IAV (Denton et al., 2019; Tan et al., 2019).

The IAV surface glycoprotein hemagglutinin (HA) is the immunodominant target of B cell responses to IAV infection and immunization (Altman et al., 2015; Angeletti and Yewdell, 2018). Pioneering studies performed over three decades ago examined the diversity of mouse B cell responses to IAV via sequencing B cell receptors (BCR) from hybridomas generated from B cells recovered at various times p.i. with the A/PR/8 (H1N1) mouse adapted strain (Caton et al., 1991; Kavaler et al., 1991; Kavaler et al., 1990). Of note, with hybridomas is impossible to discern the cell type originating the fusions. Nevertheless, comprehensive studies assessing the link between transcriptional status and clonal diversity of B cell populations at different developmental stages within or between organs after respiratory viral infections are lacking. Deciphering how BCR characteristics are linked to cell differentiation is crucial for our ability to understand and ultimately manipulate B cell responses with more effective vaccines or adjvants.

Few studies identified lung Bmem as critical in preventing IAV reinfection (Allie et al., 2019; Onodera et al., 2012). These tissue resident Bmem (Allie et al., 2019) appear to have a broader specificity compared to splenic Bmem (Adachi et al., 2015). However, virtually nothing is known about their overall transcriptional programming, their BCR profile, and whether they originate from lung-iBALT *vs.* other lymphoid organs. Better appreciation of the origin and formation of lung resident memory cells after infection is crucial first step in developing mucosal vaccines against respiratory viruses.

Germinal centers (GC) form as a consequence of rapid clonal proliferation during T cell dependent B cell responses and are the site of B cell affinity maturation through selection of high affinity clones generated via somatic hypermutation (SHM) (Mesin et al., 2016). GC entry, exit and dynamics have primarily been studied during responses to simple model antigens (Victora and Nussenzweig, 2012). A recent study suggested that following protein immunization, naive B cells with high avidity BCRs tend to immediately differentiate into PB or IgM Bmem, while B cells carrying lower avidity BCRs enter the GC (Pape et al., 2018).

Activated B cells initially acquire a light zone (LZ) phenotype, then cycle into the dark zone (DZ) where they proliferate and acquire SHM. After returning to LZ, BCR avidity is assessed by interaction with antigen on follicular dendritic cells, and current models suggest various B cell fates are determined by signals from T follicular helper cells (Mesin et al., 2016). Such fates include re-entering the DZ for additional rounds of SHM, differentiating to PB or Bmem or undergoing apoptosis. Signals that regulate terminal B cell differentiation to PB and Bmem have primarily been studied using model antigens and transgenic mice (Krautler et al., 2017; Phan et al., 2006; Shinnakasu et al., 2016; Smith et al., 1997; Suan et al., 2017a; Suan et al., 2017b; Weisel et al., 2016). The general consensus is that higher avidity B cells will differentiate into PB, while B cells of lower avidity become Bmem (Suan et al., 2017b). This was demonstrated in a study using the model hapten nitrophenol (NP) as antigen, where LZ GC B cells highly expressing Bach2 were of lower avidity and destined for Bmem differentiation (Shinnakasu et al., 2016). A subsequent study using the switched-HEL (hen-egg lysozyme) transgenic mouse model, supported this and further indicated CCR6 as marker for lower avidity LZ GC B cells becoming Bmem (Suan et al., 2017a). Interestingly, the latter study reported a high avidity Bmem subset as well. Importantly, unlike most natural responses, both NP and HEL models require only a single mutation for the germline V region in the BCR to mature from low to high avidity. Whether a similar selection of lower avidity GC cells into the memory compartment occurs after viral infection and how the selection differs between organs is unknown. This is particularly relevant to understand how the first encounter with a virus shapes Bmem formation, a central feature of the original antigenic sin phenomenon in anti-IAV responses (Henry et al., 2018; Yewdell and Santos, 2020).

To address these questions, and overcome limitations of previous studies, we have sequenced single HA-specific B cells to correlate their transcriptome with their paired heavy and light chain BCR within different organs and across time points post intranasal IAV infection. Our data provide a comprehensive resource to trace B cell differentiation upon respiratory viral infection.

## RESULTS

### scRNA-seq of antigen specific B cells after influenza infection identifies a range of B cell differentiation stages

Intranasal (i.n.) mouse infection with IAV is a well-established acute respiratory viral infection model. We infected mice i.n. with IAV PR8 and tracked antigen-specific responses at 7, 14 and 28 days post infection (dpi) by sorting antigen experienced, HA+ IgD- B cells from individual lungs, spleen and mediastinal lymph nodes (mln) (Fig 1A-B). As a control, we sorted total B cells (Live, CD3-, B220+) from spleen and lungs of two mice (Fig S1A). We subjected antigen-specific cells to single cell RNA sequencing (scRNAseq) paired with single cell B cell receptor profiling (scBCRseq). In total, we analyzed results from 8,722 cells from 2 naïve mice and 30,242 cells from 8 infected mice (1,878, 15,428, and 12,936 cells, respectively, from 7, 14, and 28 dpi).

**Fig 1.**
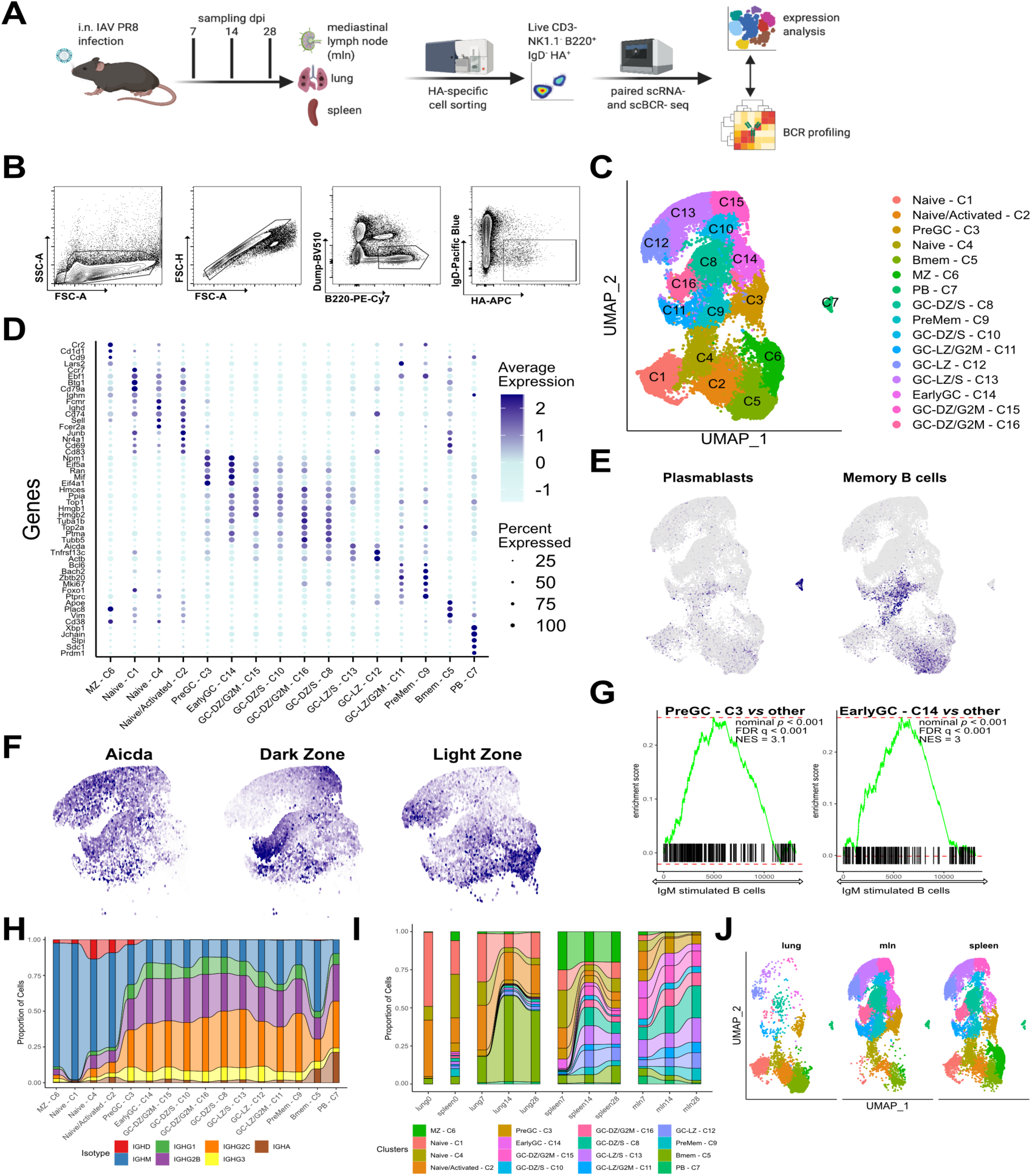
Dynamics of antiviral B cell response at single cell resolution is organ specific. A) Schematic diagram of experimental setup of influenza infection, cell sorting followed by scRNA-seq and BCR profiling. B) Representative gating for the cell sorting of single HA+ IgD- B cells. C) UMAP plot of unsupervised clustering of HA-specific B cells, combining all organs and dpi. D) Mean expression of the top 5 marker genes for each cell cluster. Color intensity denotes average expression while dot size the percent of cells expressing the gene. E) UMAP plot as in C showing average expression of gene signatures associated with plasma blasts and memory B cell programs. F) UMAP plot of GC clusters showing average expression of Aicda and gene signatures associated with dark and light zone programs. G) Enrichment score from GSEA comparing PreGC-C3 and EarlyGC-C14 to all others for genes involved in B cell activation and differentiation. H) Alluvial plot showing proportion of cells with defined antibody isotype for each cluster. I) Alluvial plot showing proportion of cells for each UMAP cluster as in C, divided by organ and dpi. J) UMAP plot of infected mice divided by organ.

Unsupervised clustering, using the Sauron implementation of Seurat package, distinguished 16 populations of HA-specific B cells that clustered according to their transcriptional profile (Fig 1C). Differential gene expression analysis allowed us to define specific cell populations (Fig 1D): naïve and activated cells (clusters C1, C4 and C2, both IgM+ and IgD+), marginal zone (MZ) B cells (cluster C6) and GC cells (clusters C3, C8, C9-16). PB (cluster C7) and Bmem (cluster C5) were confirmed by previously described gene signatures (Fig. 1E) (Bhattacharya et al., 2007). Interestingly, GC cluster C9 showed high expression of genes associated with Bmem fate (see below for further discussion). Likewise, we could readily divide GC clusters into LZ and DZ cells using sets of genes known to distinguish these cells (Fig. 1F) (Victora et al., 2010). Despite having regressed out cell cycle influence, GC clusters still separated based on cell cycle phase, as expected (Fig S1B). We identified two LZ clusters (C3 and C14) that expressed gene signatures typical for strong interactions with T follicular helper (Tfh) cells and GSEA confirmed signatures consistent with recently antigen-activated B cells (Fig 1G) (Busconi et al., 2007).

Indeed, cluster C3 cells were mostly in G1 (PreGC) while C14 cells were entering cell cycle (EarlyGC). Clusters C15, C10, C16, and C8 had similar signatures, indicative of DZ GC B cells, with only differences in cell cycle status, with C16 in G2M, C8 and C10 in S phase and C15 between G2M and G1, indicating cells exiting the cell cycle. Cluster C12 had a strong LZ signature, as did C11, which was split between G2M and G1 and C13, in S phase (Fig S1A-B).

To incorporate BCR sequence data into the overall analysis we compiled the sequence data, isotype and somatic mutations for the heavy chain sequences using the Immcantation pipeline (Gupta et al., 2015; Vander Heiden et al., 2014) and scRepertoire to define clonal status and expansion (Borcherding et al., 2020). Consistent with the recent demonstration of pre-GC class switching (Roco et al., 2019), cluster PreGC-C3 exhibited over 60% class switched Ig (Fig. 1H). Therefore, we identified PreGC-C3 cells as those receiving positive T cell help signals and that were actively recruited into the EarlyGC-C14. Supporting this hypothesis, the fraction of B cells in these two clusters was almost double at d7 compared to d14 and 28. The continued presence of these clusters weeks after infection is consistent with continued replenishment of the GC reaction (Fig. 1I). Bmem cells (cluster C5) were ∼50/50 IgM+/class switched, while all GC clusters’ cells were dominated by IgG2b/c. IgA isotype was highly enriched both in Bmem as well as in PB in all organs, suggesting a potential preferential recruitment of these cells into the effector arm (Fig. 1H and Fig S1C).

### Antigen specific clusters are organ specific and independent of day post infection

Whereas most clusters were found in all organs, surprisingly, HA+ B cell clusters were largely organ- and not dpi-specific (Fig. 1I-J). While the fraction of naïve and early activated cells was somewhat higher at 7 dpi in all organs, the most evident difference was the distribution between cell types in the three organs studied. The mln is characterized by strong GC activity with a smaller proportion of Bmem (3-6%) and a considerable number of PB (from 7% at day 7 to 2% at d28) (Fig. 1I). Conversely, we could detect very low HA+ GC B cells in the lungs (∼10%) but a remarkably high number of Bmem that rose from 10% at day 7 to ∼30% of HA+ cells at d14 and 28. PB were constant between 1-2%. HA+ B cells in the spleen exhibited strong GC activity and relatively constant proportion of Bmem (8-13%) and PB (0.5-1%). Interestingly, PB proportion was the highest in mln at day 7 but lowest in other organs. This, linked to mutation analysis (Fig 6) showing nearly germline BCRs in PB, indicates an early and selective expansion of PB in the mln. Coordinately, we detected a burst of Bmem in mln at d7.

Overall, these data demonstrate early expansion and differentiation of HA-specific cells to PB and Bmem in the mln, and persistence of Bmem in lungs.

### Identification of Bmem precursors using scRNA-seq

The dynamics of GC at single cell resolution after viral infection have not been previously elucidated and, further, the identity of Bmem precursors in the GC is controversial (Laidlaw et al., 2017; Shinnakasu et al., 2016; Suan et al., 2017a). In order to address these issues and decipher the pattern of Bmem differentiation, at the single cell level, we performed trajectory analysis using Slingshot (Street et al., 2018) and RNA velocity analysis with scVelo (Bergen et al., 2019). RNA velocity analysis suggests that PreGC-C3 differentiate to EarlyGC-C14 and subsequently enter GC (Fig 2A), in accordance with GSEA (Fig. 1G). Further, velocity analysis suggests that PreMem-C9 could potentially have some backflow to Bmem (C5). For trajectory analysis with Slingshot we removed cluster PB-C7, as the cluster was clearly disconnected form all others. the trajectory analysis, with preset start at PreGC-C3, showed a major trajectory going from LZ to DZ and to LZ again, with cells exiting from PreMem-C9 and differentiating to Bmem-C5 (Fig. 2B). Changes of gene expression along Slingshot pseudotime showed a marked switch in transcriptional programming at PreMem-C9 (Fig 2C). PreMem-C9 cluster is unique in that cells highly express several GC marker genes (*Mki67, Aicda, Bcl6*) as well as *Foxo1, Bach2, Cd22* and have high mitochondrial content (Fig 2D). All these genes and features have been implicated in Bmem development (Chappell et al., 2017; Jang et al., 2015; Shinnakasu et al., 2016). Altogether, this cell population closely resembles the Bmem-precursor cell population previously identified by Shinnakasu *et al*. (Shinnakasu et al., 2016).

**Fig 2.**
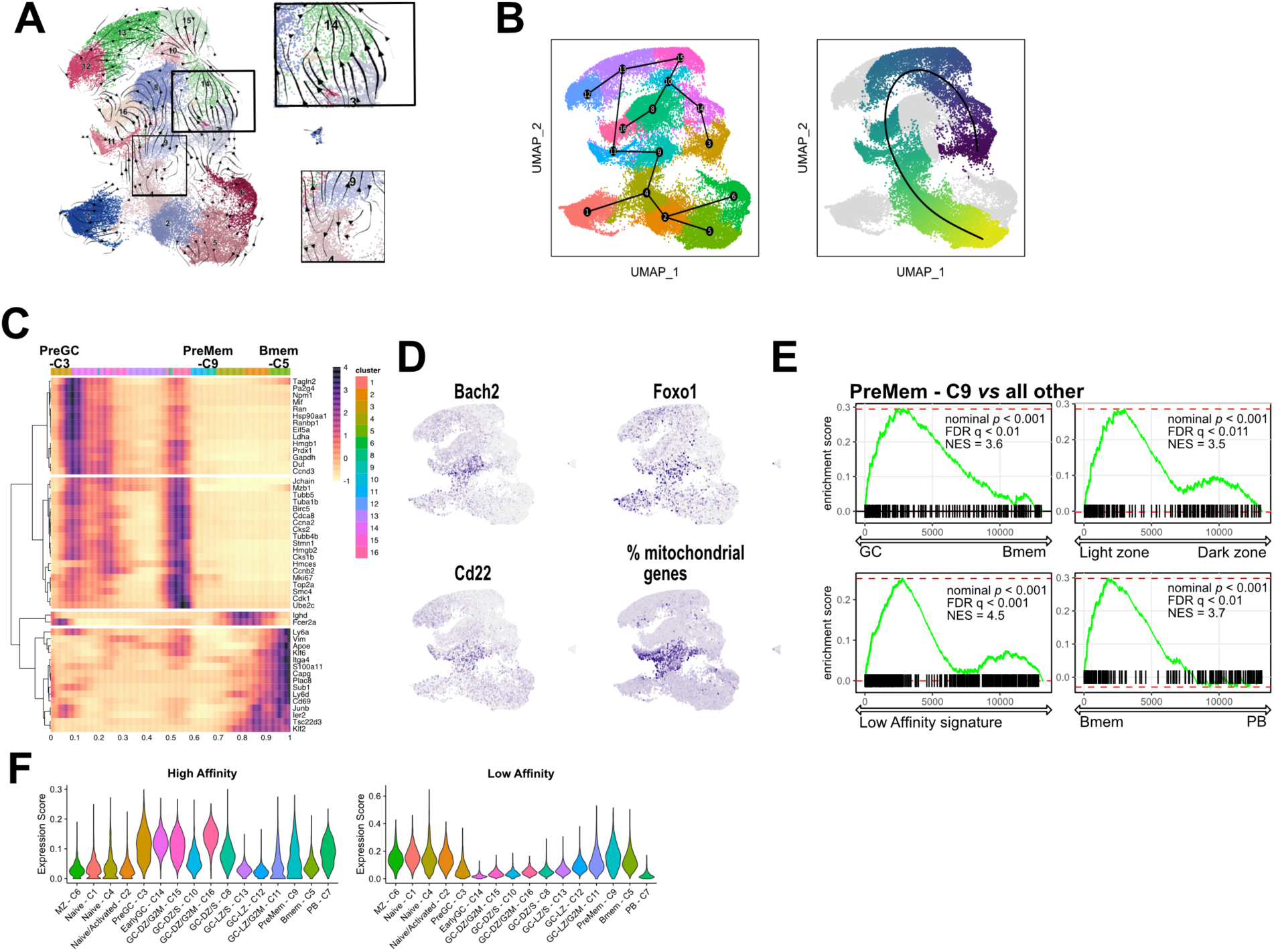
RNA velocity and trajectory analysis identify cluster 9 as memory B cell precursors. A) RNA velocity determined by scVelo projected on UMAP. Arrowheads determine predicted direction of the cell movement and arrow size determines strength of predicted directionality. In the squares are highlighted cells moving from PreGC-C3 to EarlyGC-C14 (top) and cells moving from PreMem-C9 (bottom). B) Trajectory inference by Slingshot projected on UMAP with PreGC-C3 selected as starting cluster. On the right the same graph with pseudotime coloring. Cluster PB-C7 was excluded as clearly disconnected from the others. C) List of differentially expressed genes over trajectory-based pseudotime. Colors on top indicate clusters. D) UMAP plot of GC clusters showing average expression of selected genes. E) Enrichment score from GSEA comparing PreMem-C9 to all others for genes involved in GC program, LZ program, memory cell program and low affinity signature as described by Shinnakasu et al. (2016) F) Violin Plot showing High and Low affinity gene expression scores by UMAP clusters as defined by Shinnakasu et al. (2016)

To further confirm that PreMem-C9 was the Bmem precursors cluster, we ran a series of GSEA analyses. We defined high and low affinity score based on average expression of genes corresponding to high and low affinity cells in Shinnakasu et al (Shinnakasu et al., 2016) at the single cell level. GSEA analysis identified PreMem-C9 as a LZ GC B cell population enriched in the low affinity score signature as well as more similar to Bmem than PB (Fig 2E). When assigning low and high affinty scores we confirmed PreMem-C9 as the highest among GC clusters for this signature. Likewise, the Bmem population was enriched for the low affinity signature. Conversely, most of the other GC clusters were enriched for the high affinity signature (Fig 2F). Thus, cluster PreMem-C9 likely represents a Bmem precursor population in the GC, characterized by high *Mki67, Bcl6, Cd22, Bach2, Foxo1* expression and high mitochondrial content.

### Bmem in the lungs have a distinct transcriptional profile, independent of isotype

Having found a different proportion of Bmem in different organs (Fig 1I-J) we investigated whether their transcriptional programming varied with anatomic location. Unsupervised analysis of the Bmem population (C5) only, revealed 8 subclusters, with the main determinant of separation being lung *vs.* non-lung localization (Fig 3A-D). Clusters 0, 1 and 4 were almost exclusively made up from lung Bmem (Fig 3C). These cells were of different isotypes and were both mutated and unmutated, thus probably of GC-dependent and independent origins (Fig 3D). While the number and proportion of IgA Bmem in the lung was higher compared to spleen(Fig S2A), B cell heavy chain class was only a partial determinant in the cell segregation. IgM+ Bmem were the principal determinant of Bmem-cluster 2 (Fig 3B), but were present in other clusters.

**Fig 3.**
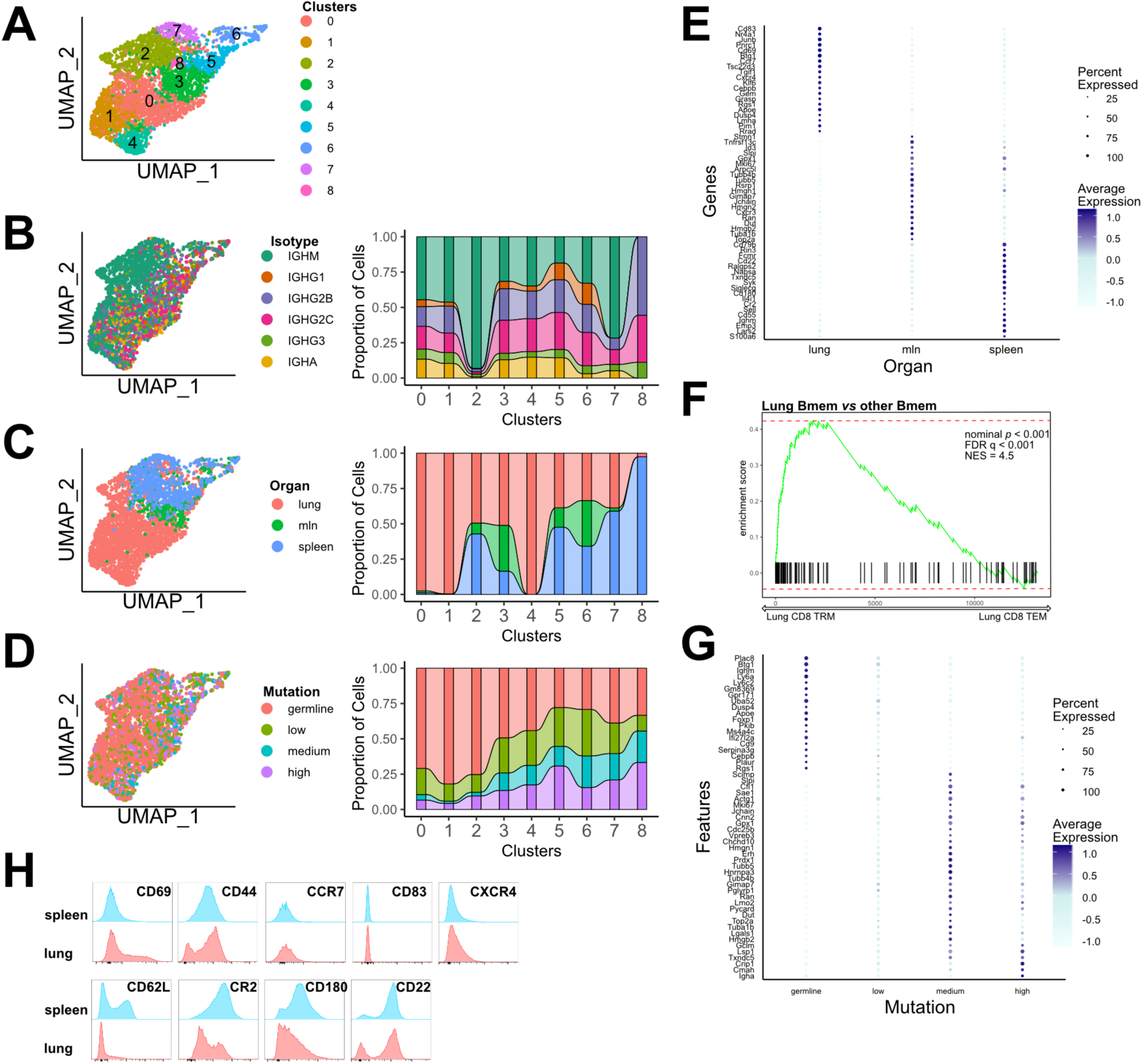
Memory B cells in the lungs have a distinct transcriptional programming compared to spleen and mln. A) UMAP plot of unbiased clustering of HA-specific memory B cells (C5 in Fig 1C), combining all organs and dpi. B) UMAP plot of unbiased clustering of HA-specific memory B cells as in A, colored by BCR isotype. On the right alluvial plot showing proportion of cells with defined isotype per cluster. C) UMAP plot of unbiased clustering of HA-specific memory B cells as in A, colored by organ. On the right alluvial plot showing proportion of cells, belonging to a specific organ per cluster. D) UMAP plot of unbiased clustering of HA-specific memory B cells as in A, colored by BCR mutation rate. Germline (not mutated), low (up to 1% nucleotide mutation), medium (up to 2%) and high (over 2% mutation). On the right alluvial plot showing proportion of cells with defined mutation rate per cluster. E) Mean expression of the top 20 marker genes for each organ for Bmem. Color intensity denotes average expression while dot size the percent of cells expressing the gene. F) Enrichment score from GSEA comparing lung Bmem to all others for genes expressed by CD8 TRM G) Mean expression of the top 20 marker genes for cells divided by mutation rate for Bmem. Color intensity denotes average expression while dot size the percent of cells expressing the gene. H) Flow cytometry histograms showing expression of indicated genes by memory B cells (Dump- B220+ CD38+ IgD- IgM-) in lungs and spleen. Representative experiment of 3 biological replicates with 3 mice each.

Rather, several genes strongly contributed to differential clustering between lung and spleen/mln clustering (Fig 3E). Interestingly, among the most differentially upregulated gene in the lungs were *Cd69*, an adhesion molecule linked to tissue residency of immune cells, together with *Cd83, Ahr, Ccr7, Cxcr4* and *Cd44*. Conversely spleen and mln had significantly higher *Sell* expression (encoding CD62L), together with *Cd22*, Cr2, *Bcl2* and *Cd55* (Fig 3E). GSEA analysis on CD8 tissue resident memory signatures revealed striking similarity of lung Bmem to CD8 tissue-resident memory (TRM), as opposite to spleen and mln Bmem (Fig 3F) (Mackay et al., 2013).

Validating the observed differences, PB and GC transcriptional profiles appeared largely similar between organs (Fig S2D-K). The only detected difference was between IgA PB and others, as previously reported (Fig S2E) (Neu et al., 2019; Price et al., 2019). Thanks to the power of our approach, we detected transcriptional differences in Bmem and PB between germline (GC-independent) and mutated (GC-dependent) cells, suggesting long term functional differences, depending on cell origin (Fig 3G and Fig S2B and Fig S2L). Germline (and probably GC-independent) Bmem expressed higher levels of *Btg1, Foxp1, Plac8*, and other genes that control cell proliferation and differentiation. On the other hand, highly mutated Bmem (probably GC-dependent) expressed higher levels of *Jchain* (IgA specific), *Slpi, Txndc5, Cmah* and others associated with programming towards PB differentiation

To validate the transcriptional differences detected in Bmem, we infected mice and analyzed lung and spleen B cells at 14dpi by flow cytometry. This revealed upregulation of CD69 and CD44 in lung Bmem consistent with scRNAseq data. CD83 and CXCR4, even if upregulated at the mRNA level, were not detected on the Bmem surface, as expected given their role in GC. Splenic Bmem had higher CD62L, CR2 and CD22 expression consistent with scRNAseq data (Fig 3H).

Collectively, these data demonstrate a distinct transcriptional profile of Bmem in the lung with hallmark of activation and tissue residency. The low number of HA+ GC B cells in the lung suggest that the majority of lung Bmem are either GC independent or derived from GC in other organs and acquiring a new transcriptional signature once they take residence in the lungs.

### Vh gene usage is mainly organ specific but shared among cell types

We assigned V gene usage using the Immcantation pipeline according to IMGT standard (Giudicelli et al., 2011). As expected, the unselected naïve repertoire was composed of a large number of Vh genes while we could start observing selection already at day 7 after infection (Fig 4A). At 7dpi, Vh1-63 expressing B cells (Cb site specific) dominated the PB and GC response on D7 (Fig S3A), as previously reported (Angeletti et al., 2017; Kavaler et al., 1990; Rothaeusler and Baumgarth, 2010), producing germline or near germline Abs (Fig S3B). Extending previous findings, we found an increased proportion of these B cells also within the Bmem compartment in unmutated form, indicating that these B cells, with high avidity germline BCR not only dominate the extrafollicular PB response but also differentiate into IgM and switched Bmem (Fig S3C). At day 14 with all individual mice, Vh gene usage became far more polarized, indicating vigorous clonal selection. Critically, by d28 one Vh family dominated in each mouse, ranging from 35% to 62% of the response (Fig 4A).

**Fig 4.**
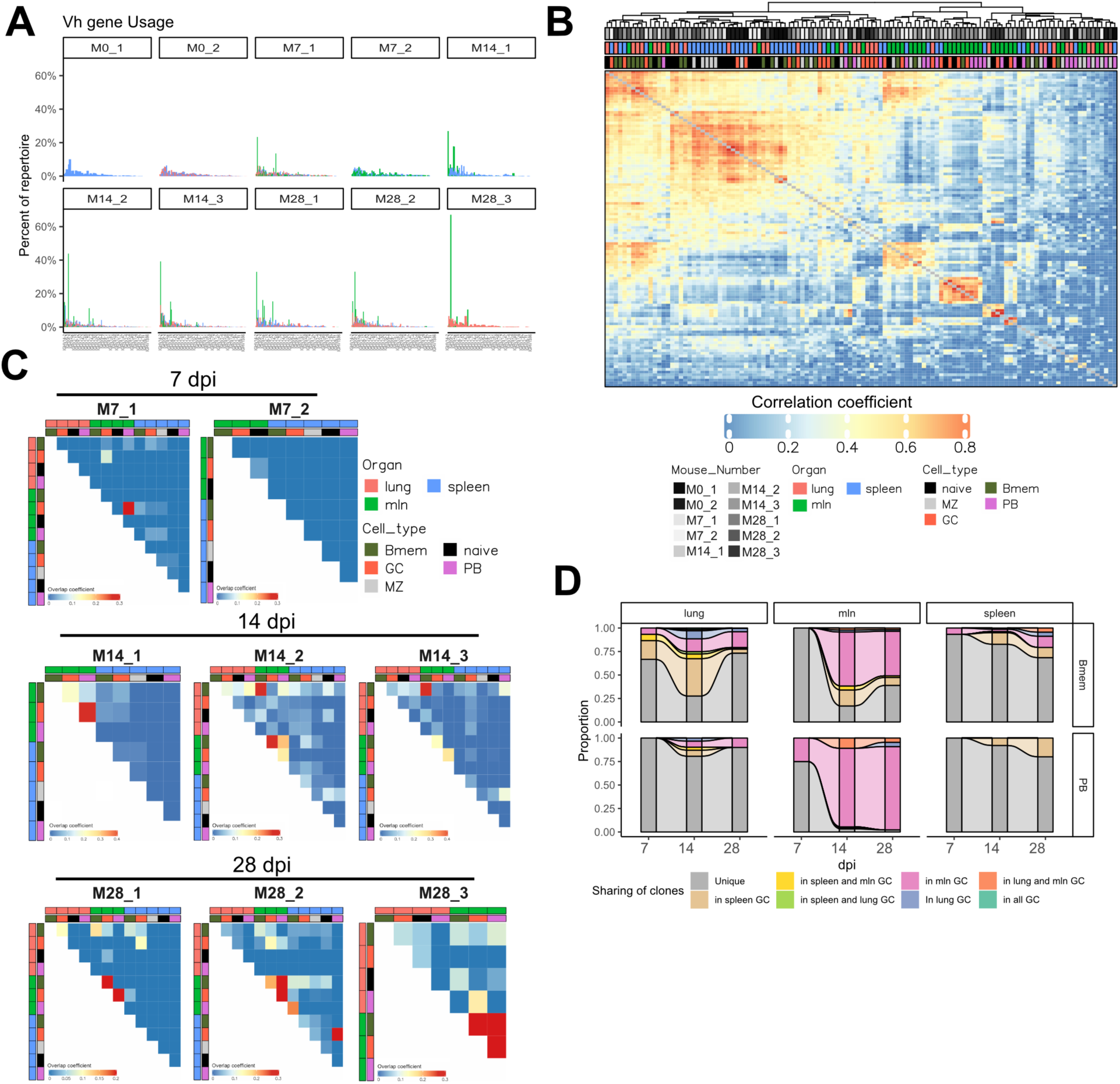
Memory cells broadly disseminate in several organs. A) Percentage of cells using a specific Vh gene for each mouse, divided by organ. B) Hierarchical clustering of Pearson’s correlation of the V gene repertoire. Each tile represents the correlation of V gene repertoire. Color intensity indicates correlation strength. See Fig S4 for p values. C) Overlap between B cell clones in different organ and cell types, divided by mouse. Each tile represents the overlap coefficient of clones. Color intensity indicates overlap strength. D) Alluvial plot showing clonal origin of Bmem and PB based on CDR3 sequence with germline (GC-independent) cells excluded from analysis. Top row show Bmem and bottom row PB, divided by organ at each dpi. Grey bar indicates that clones were not found in any GC, while color indicates that clonal relatives were found in GC.

When comparing cell clusters at different time points our data shows some Vh genes undergo selected as early as 7dpi in GCs, and dominant clones also appear in PB at 14dpi (Fig S3D). Finally, we detect some skewed Vh usage in the Bmem population at 28dpi, consistent with prolonged GC selection (Fig S3D). Selection of Vh, D and Jh, genes was mainly cell type rather than organ specific (Fig S3E-F). V gene usage was private to individual mice, except for three genes that were consistently overrepresented in all mice (Vh14-2, Vh1-81 and Vh1-63) (Fig S4A).

We then analyzed the Vh gene usage overlap between different mice, organs and cell types, as defined by UMAP clusters, using a Pearson’s correlation matrix (Fig 4B and Fig S4B) (Greiff et al., 2017a). This analysis gives information on both overall Vh gene usage, selection and clonal expansion. This revealed that 1) the V gene repertoire is most clustered by the high similarity between lung and spleen B cells, 2) numerous Vh genes can be used to generate HA binding Abs, 3) most of them are selected into GC and Bmem compartments, 4) selection in to PB compartment is limited to much fewer clonotypes, which are unusually excluded from GC and Bmem compartments.

Analyzing clonal overlap based on both heavy and light chain (Fig S4C), revealed that clustering is dominated by individual mice, extending many prior findings (Greiff et al., 2017a; Greiff et al., 2015; Greiff et al., 2017b; Miho et al., 2019) that even in mice with nearly identical genetic backgrounds most B cells generate repertoires that emerge stochastically.

### Memory B cells disseminate in several organs

The CDR3 Pearson’s correlation between individual mice, organs and cell types indicates that clonal overlap is mostly organ specific with notable exceptions. In several mice, the Bmem populations in lungs were overlapping with GC and Bmem populations in mln, rather than other B cell populations in lungs themselves. PB in mln were strongly correlated with mln GC in most mice but the same wasn’t always true for PB in spleen and lung (Fig 4C and Fig S4D).

We further investigated clone sharing between PB, Bmem and GC in different organs. We only considered mutated, likely GC-dependent clones. By day 14 we could assign almost all PB in mln to have GC origin by clonal relationship (Fig 4D). This was not true for most of the lung and spleen, due at least in part to the low degree of clonal expansion and limited sampling. However, it should be noted that, at least for the spleen and mln, overall diversity for each cluster was similar (Fig S5). For Bmem in spleen we idenfied clonal relatives for only ∼20% of the cells. This could be due to smaller clonal families that make sampling limiting. We could, however, track as high as 75% of lung and mln Bmem at day 14 (Fig 1H). Surprisingly, we observed sharing of GC-derived Bmem, with spleen GC being a source of Bmem in mln and lungs and mln-derived Bmem present in the lungs and spleen. These data are consistent with a high degree of dissemination of GC-derived Bmem between organs.

### Clonal expansion is organ specific and highly expanded clones seed both PB and Bmem compartments

Next, we examined clonal expansion. We found that up to 75% of mln antigen specific B cells belonged to expanded clonotypes. By contrast, at most 20% of B cells in the lung/spleen were expanded (Fig 5A). Consistently, the top 50 expanded clones represent more than 50% of the repertoire in mln, while just 15-30% in lungs and spleen (Fig S6A). Expectedly, we detected no sign of clonal expansion in naïve mice. Clonal expansion by cluster and dpi shows lower expansion in splenic GC B cells compared to mln B cells (Fig 5B). Clonal expansion of the various GC clusters was uniform.

**Fig 5.**
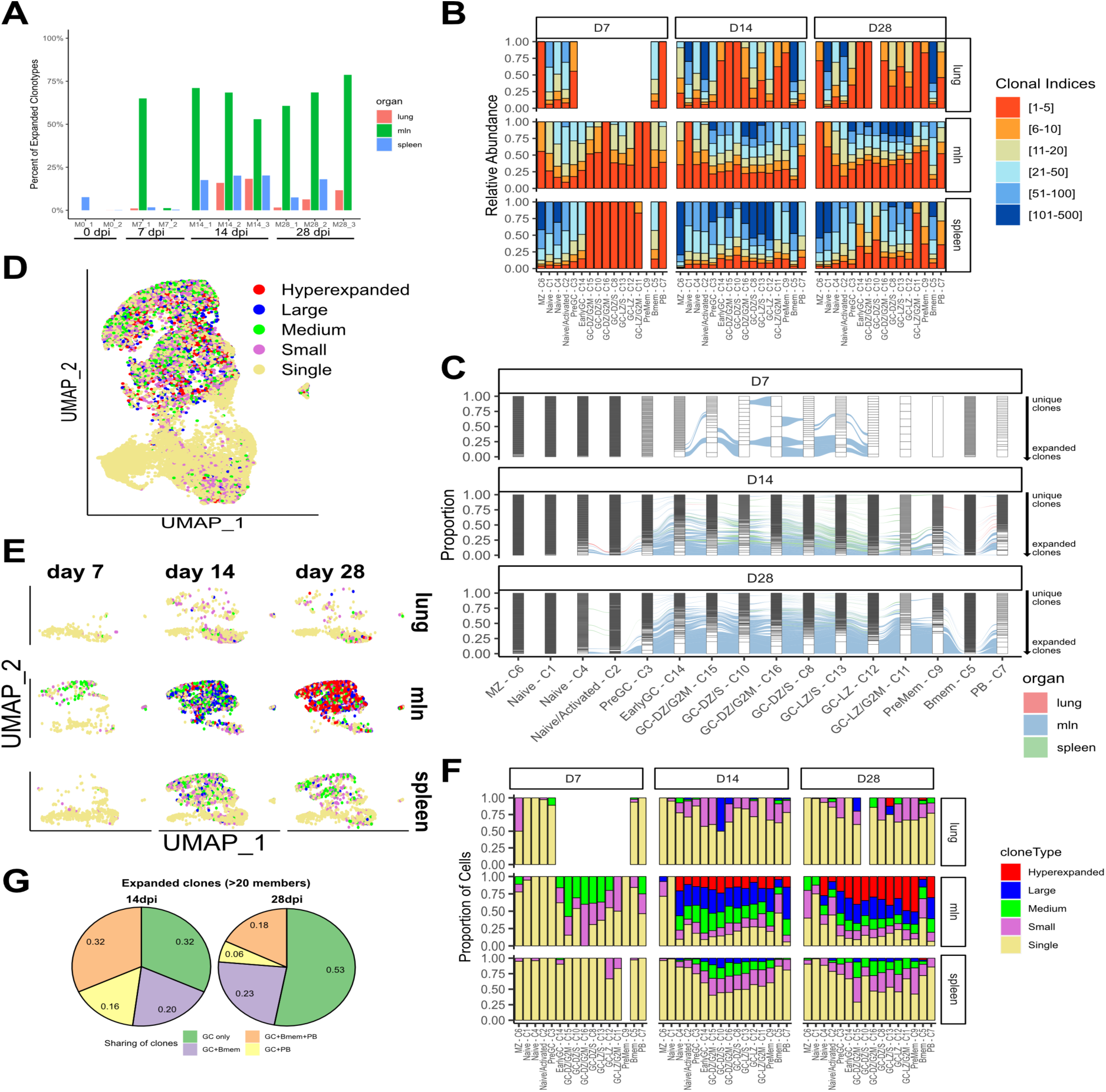
Clonal expansion in GC is organ specific. A) Graph showing the percentage of expanded clonotypes for each mouse, divided by organ. B) Graph showing clonal expansion for each cluster, divided by dpi and organ. Clones are order by their abundance for each cluster, dpi and organ and color indicate the repertoire space occupied by the top X clones as shown in figure legend. C) Alluvial plot showing clonal sharing between clusters and organs at different dpi. Each cluster is divided in several bars, representing individual clones, and the height of each represents the proportion of the cluster occupied by that clone. Connecting lines indicate sharing of clone with colors indicating the organ. D) UMAP plot of infected cells for all organs and dpi, colored by clonal expansion status. Clones were defined as single (1 cell), small (between 1 to 5 cells), medium (between 6 to 20 cells), large (between 21 to 100 cells) and hyperexpanded (over 101 cells) clones. E) UMAP plot of infected cells divided by organ and dpi, colored by clonal expansion status. F) Graph showing proportion of cells for each clonal expansion status for each cluster, divided by dpi and organ. G) Pie chart showing distribution of expanded clones (more than 20 cells sequenced) in different clusters. Numbers in the chart indicate frequency.

Bmem are the most BCR diverse compartment, and had higher numbers of unique clonotypes (Fig S5). Conversely, PB started with few highly expanded clonotypes at day 7, became more diverse by day 14 and narrowed again over the next two weeks. We generated alluvial graphs to assess the extent to which highly expanded clonotypes are shared between clusters or preferentially expanded in certain clusters over time (Fig 5C). At day 7 only a few clonotypes in the GC clusters were shared among multiple clusters, regardless of clonal expansion state. By day 14 about 30-50% of the highly expanded clones were present in all GC clusters and most of the highly expanded clones populate Bmem and PB subsets. Clonal sharing between GC and Bmem and PB was independent of clonal size. At day 28, more than 50% of BCR sequences were shared between GC clusters, in particular among highly expanded clones. Consistent with clonal expansion state, about 50% of PB derived from highly expanded clones. As in previous dpi, Bmem were originated not only from highly expanded GC families but also from smaller clones (Fig 5C). When looking at individual mice, M14_2 and M14_3 at d14 stood out as that most of the clonal sharing was only between splenic GC and not mln GC, while M28_3 at d28 had more than 75% clonally expanded clones shared between clusters (Fig S6B).

We visualized clonal expansion by arbitrarily dividing clones into five categories: single (1 cell), small (between 1 to 5 cells), medium (between 6 to 20 cells), large (between 21 to 100 cells) and hyperexpanded (over 101 cells) clones and rendered them on the UMAP plot (Fig. 5D). The UMAP clearly shows that the majority of expanded cell clones are in the GC clusters, but also in PB and Bmem clusters. As expected, cells in naïve and MZ clusters mostly belonged to single clones. Interestingly, cells in PreGC-C3 cluster, which we propose to represent cells just entering the GC and undergoing Tfh interactions, were mostly unique, consistent with our hypothesis.

Surprisingly, splitting the UMAP according to organ and day (Fig. 5E-F and Fig S6C) reveals that GC clonal expansion is organ dependent. In mln, medium expanded clones are already present at d7 p.i. and large and hyperexpanded clones by d14. Conversely, splenic GC had few medium sized clones at day 14 and maintained an essentially unmutated expansion profile at day 28. Likewise, GC clones in the lungs were mostly single or small with occasional medium sized clones appearing. These findings are quite surprising, as they clearly show organ-dependent regulation of GC clonal expansion. Notably, the number of analyzed GC cells in mln and spleen at day 14 is nearly identical (n= 2125 vs 1828). Nevertheless, to verify that sampling differences didn’t affect our day 28 observation, we randomly downsampled mln d28 to the same number of GC cells of spleen and reassigned the expansion status of clonotypes (Fig S6D). We still detected the majority of GC B cells to be either large or hyperexpanded, in stark contrast with spleen GC, validating our conclusion.

Focusing on highly expanded clones (more than 20 cells), we could define four patterns: clonotypes present in GC only, GC and PB, GC and Bmem and GC, Bmem and PB. Remarkably, the proportion of highly expanded clones found only in the GC increased from 33% to about 50% from day 14 to 28. Conversely, the fraction of GC clones shared only with PB decreased, while the fraction of Bmem derived from GC stayed constant by 28dpi (Fig 5G). Together with overall clonal expansion status, this observation indicates that a constant number of GC derived Bmem are generated as the immune response progresses while the PB diversity decreases as PB clones expand.

### GC-derived PB and Bmem have similar mutation rate and avidity for antigen

Finally, we investigated how BCR somatic mutation status relates to cell type and clonal expansion of HA-specific B cells.

In line with what would be expected, naïve and marginal zone cells had almost no mutations, while GC cells were overall the most mutated followed by PB and Bmem (Fig 6A). Separating by dpi and differentiation clusters we found that d7 mice had very few mutations, regardless of cell type (even in PB and Bmem), indicating that PB and Bmem initially derive from expansion of unmutated cells. The mutation frequency increased at later dpi with similar trends for all cluster and organs (Fig 6B). When considering the different heavy chain classes (Fig 6C and Fig S7A), we did not detect major differences between clusters and organs with two exceptions: IgM cells were generally less mutated than class switched cells and IgA cells tended to have a higher mutation rate, starting from d14. B cell mutation frequencies between mice were comparable (Fig S7B).

**Fig 6.**
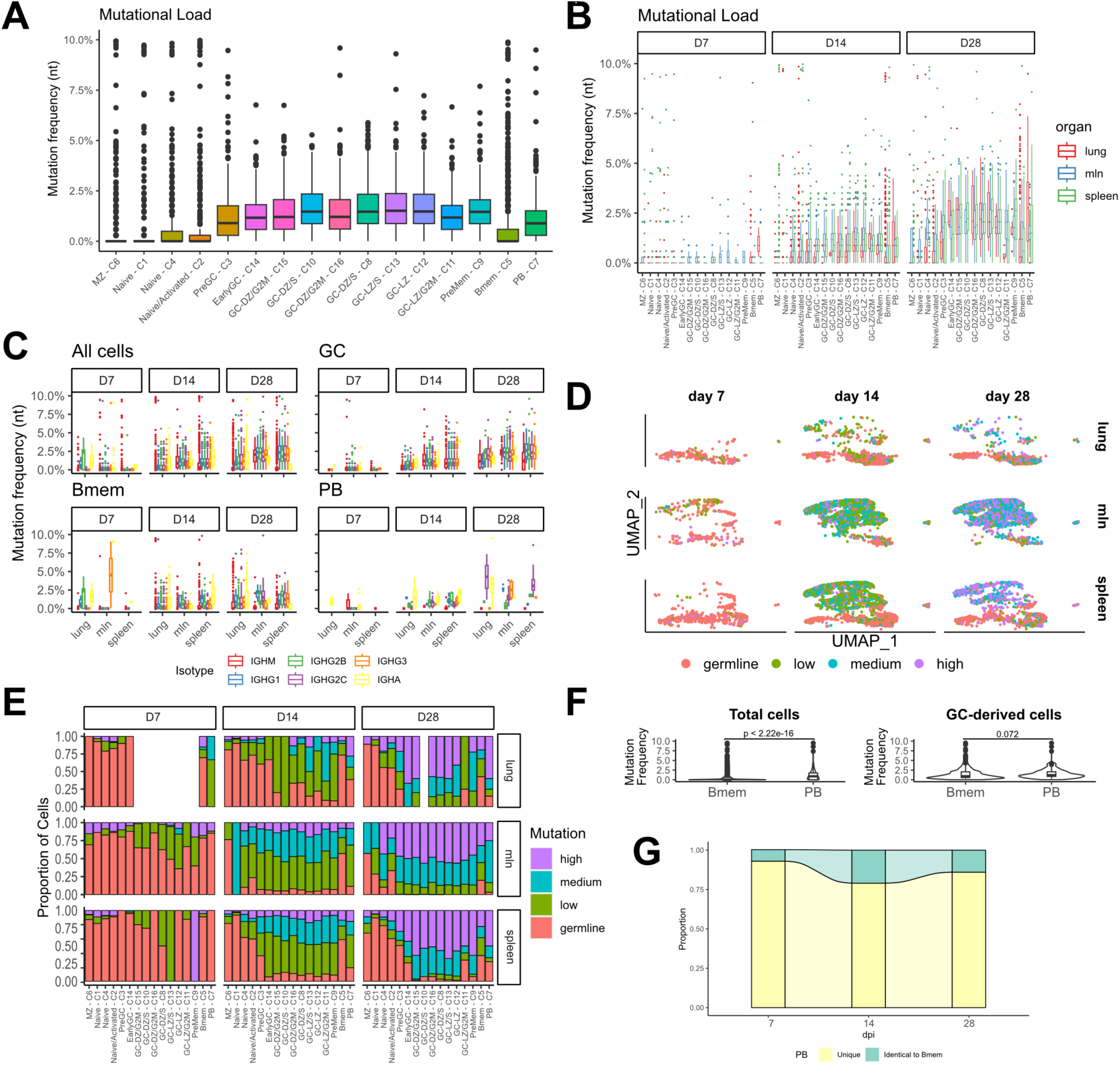
Sustained generation of highly mutated Bmem. A) Graph showing Vh gene mutation frequency divided by UMAP clusters as in 1C. B) Graph showing Vh gene mutation frequency for each cluster, divided by dpi and organ. C) Graph showing Vh gene mutation frequency for each organ, divided by dpi and isotype. Two-way ANOVA with Tukey’s post test: for “All cells” at d14 each isotype vs IgM p<0.0001, IgA vs IgG2b p<0.05, IgA vs IgG2c p<0.001, IgA vs IgG3 p<0.01, IgG2c vs IgG1 p<0.001. For “All cells” at d28 each isotype vs IgM p<0.0001, IgA vs IgG1 p<0.01, IgA vs IgG2b p<0.05, IgG2b vs IgG2c p<0.0001, IgG2c vs IgG3 p<0.01, IgG2c vs IgG1 p<0.0001, other comparisons ns. For “GC” d28 IgG1 vs IgM p<0.001, IgG2b vs IgM p<0.0001, IgG3 vs IgM p<0.01, IgG2c vs IgG1 p<0.0001, IgG2c vs IgG2b p<0.0001, IgG2c vs IgG3 p<0.0001. For “Bmem” d14 IgA vs IgM p<0.0001, IgA vs IgG2b p<0. 0001, IgA vs IgG2c p<0. 0001, IgA vs IgG3 p<0.001. For “PB” d28 IgA vs IgM p<0.0001, IgA vs IgG2b p<0. 0001, IgA vs IgG2c p<0. 0001, IgG1 vs IgM p<0.01, IgG2b vs IgM p<0.01. All other comparisons non significant. D) UMAP plot of infected cells divided by organ and dpi, colored by mutation rate. Germline (not mutated), low (up to 1% nucleotide mutation), medium (up to 2%) and high (over 2% mutation). E) Graph showing proportion of cells for each mutation rate for each cluster, divided by dpi and organ. F) Violin Plot comparing mutation frequency of total and GC-derived Bmem vs PB. Statistical differences were tested using Student’s t test. G) Alluvial plot showing proportion of PB with a sequence totally identical to a Bmem, divided by dpi.

To facilitate mutation analysis, we divided the cells in four discrete bins: germline (not mutated), low (up to 1% nucleotide mutation), medium (up to 2%) and high (over 2% mutation) (Fig 6D). By day 14 most of GC cells carried BCR with low to medium mutations while at day 28 they had medium/high mutation rate. Comparing the mutation data with clonal expansion data (Fig 5F) highlights the different dynamics between mln and other organs.

Approximately 75% of Bmem remained germline at day 14 and 50% at day 28 (Fig 6E). While we can’t determine the timing of their production, the increased proportion of highly mutated Bmem suggested recent origin. More than half of the unmutated Bmem were of IgM isotype but we also detected IgG and IgA (Fig S2C). Unexpectedly, we found that when excluding non-mutated, likely GC-independent cells, overall mutation rates of PB and Bmem BCR were statistically indistinguishable, with the exception of Bmem in the lungs and spleen at d28 having lower mutation rate than PB in the same organ (Fig 6F and Fig S7C). Similarly, PreMem-C9, identified by trajectory analysis to be Bmem precursors (Fig 2), had undistinguishable mutation rate and clonal expansion profile compared to all other clusters. In addition, we found that 22% of PB had BCR sequence identical to a Bmem. Further, mutation distribution did not correlate with clonal size, with families with only 5 members already showing members with high mutation rate (Fig S7D).

While an imperfect proxy, higher SHM usually reflects increased Ab binding avidity (Gitlin et al., 2014; Kocks and Rajewsky, 1988; Neu and Wilson, 2016). Indeed, high and low avidity signatures correlated with mutation rate (Fig S7E). As GC-derived Bmem are thought to originate from lower affinity cells as compared to PB (Shinnakasu et al., 2016; Suan et al., 2017a), we would expect GC-derived PB to have higher mutation rate as compared to Bmem but this was not the case in our infection model (Fig 6F).

The mutation data suggests only a minor contribution of affinity-based selection for GC B cells becoming Bmem vs PB. To test this hypothesis, we generated mAbs from mutated Bmem and PB that are members of large clonal families We generated clonal trees for five families (one from M1 and two from M2 at 14dpi and one each from M5 and M6 at 28dpi) that contributed differing proportions of cells to GC, Bmem and PB populations (Fig 7A). The branching point for differentiation into PB *vs.* Bmem appeared to be random. In more complex trees, some branches gave rise to both PB and Bmem.

**Fig 7.**
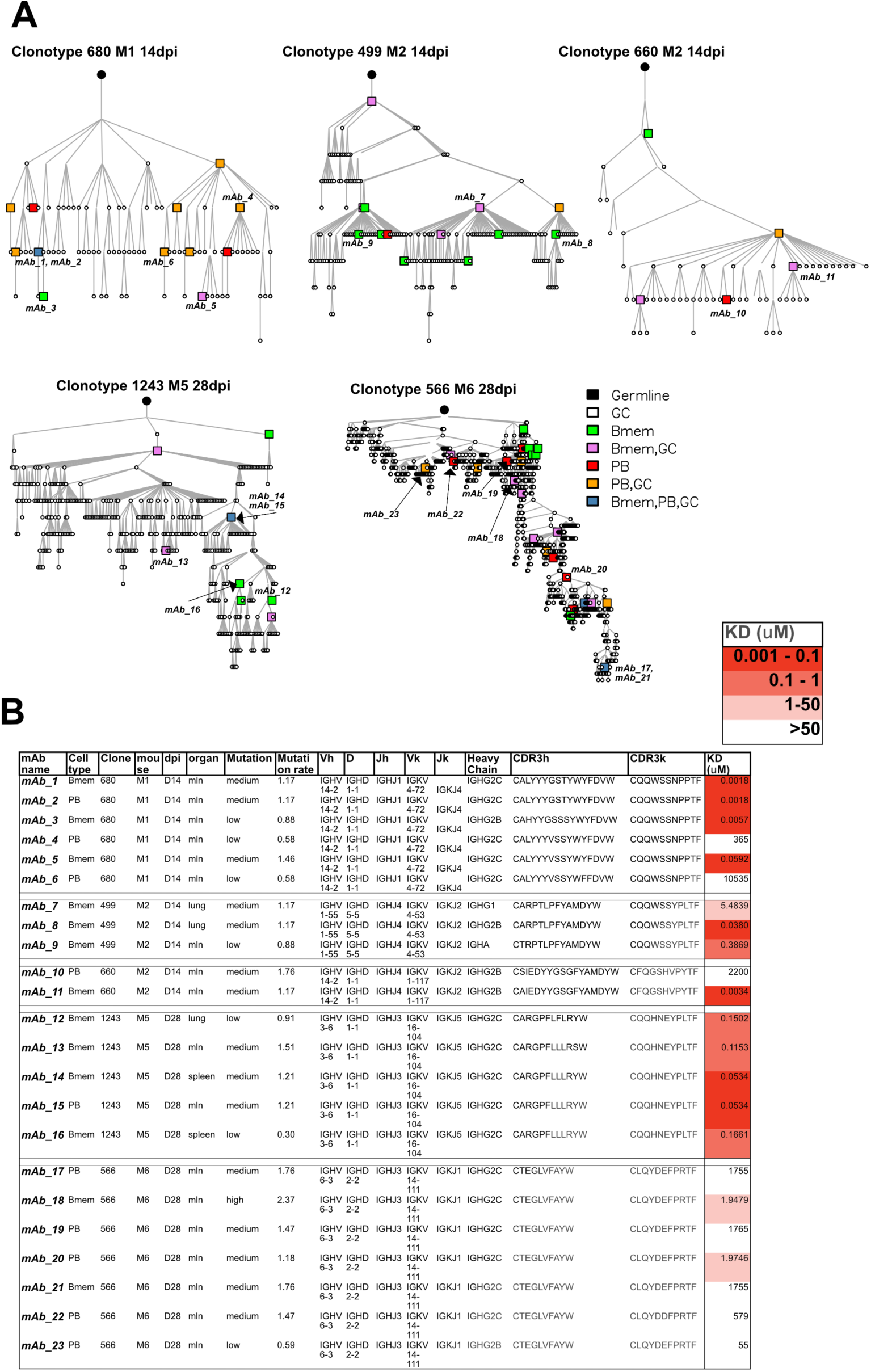
mAbs derived from Bmem and PB have similar affinity for HA. A) Clonal trees of five selected clonal families from 4 mice at different dpi. Color indicates cell types, where each circle or square is a cell that was sequenced in our experiments. Where symbols are missing between junctions, denotes an inferred member of the clonal family. Expressed mAbs are indicated by name. B) Characteristics of the expressed mAbs including KD value measured by BLI.

We expressed 23 representative switched mAbs (13 from Bmem and 10 from PB) and calculated their affinity to HA by biolayer interferometry (BLI) (Fig 7B and Fig S7F). Of note, 3 of the selected mAbs were identical between PB and Bmem. BLI affinity measurement showed no pattern of differential affinity between Bmem and PB. Indeed, the major determinant of affinity was the clonal family and not the cell type or number of mutations, similar to what was recently reported for Bmem recall after immunization (Mesin et al., 2020). Surprisingly, mAbs from the highly expanded clonotype 566, with more than 700 sequenced cells (>70% of all cells of M28_3) exhibited low to extremely low avidity for HA by BLI. In an extreme example of diverse avidity within a single clonotye, in clone 660, mAb 11 and mAb 10, which differ by 2 amino acids (with 1 in the CDR3), exhibit a nearly million-fold difference in KD (2.2 mM vs. 3.4 nM). We confirmed the BLI measurements by testing the mAb by ELISA on HA, HA from virus and PR8 virus treated at pH5 to expose hidden epitopes (Fig S7G-I). Interestingly, pH treatment affected mostly members of one clonal family (1243) by decreasing their apparent KD, while only one mAb had increased Kd upon pH treatment (mAb 10). While the results did not fully recapitulate the BLI measurements, they confirmed that there was no difference in apparent avidity between Bmem and PB between the same family, that the clonal family was the main determinant in avidity difference and that all mAb bound virus.

Overall, these data indicate that following viral infection: 1) a sizeable number of high avidity GC-dependent Bmem are generated as early as two weeks p.i. 2) GC-derived Bmem and PB possess BCR with similar number of mutations and avidity for antigen. 3) Clonal selection is not strictly based on the measured avidity of soluble Abs encoded by the clones.

## DISCUSSION

B cell responses are the cornerstone of preventing viral infections. Better understanding of how antigen specific B cell immunity develops after a respiratory viral infection is crucial for designing effective vaccines for influenza viruses, parainfluenza viruses and SARS-CoV-2. How antigen-specific B cells differentiate prior to GC and from the GC to PB and Bmem remains elusive. Here, by combining antigen-sorting, scRNA-seq and scBCR-seq we have generated a detailed map of differentiation stages of antigen specific B cells in response to respiratory viral infection. By analyzing events in lungs, draining LN, and spleen, our data elucidate the complex mechanisms involved in B cell responses to infection.

Most seminal discoveries in B cell biology have been made in mice immunized with haptens or simple, monovalent protein antigens (e.g.NP, OVA, HEL, CGG). Such models do not, however, fully recapitulate the complexity of infectious agents, each of which expresses dozens to hundreds antigens, and also idiosyncratically activates innate immunity, which sculpts the adaptive response. Indeed, using a more complex protein immunogen for immunization (IAV-HA) can challenge established principles of B cell differentiation (Kuraoka et al., 2016; Mesin et al., 2020).

Our findings clearly demonstrate the critical importance of using viral infections to study B cell responses. Following IAV lung infection, we used gene signatures to assign identities to scRNA seq clusters, and differentiate LZ to DZ in the GC. We observed that the proportion of Bmem, PB and GC HA-specific cells did not vary with time p.i. infection but was specific for each organ. This was also true for all specific GC clusters. Not surprisingly, PreGC-C3 and earlyGC-C14, which include unmutated cells interacting with Tfh and entering the GC reaction were overrepresented on day 7. Within PreGC-C3 and earlyGC-C14 we could detect progressive class switching as cells were approaching GC, consistent with recent findings of pre-GC class switching (Roco et al., 2019).

The regulation of B cell differentiation was organ specific. In particular, lungs harbor a large number of HA-specific Bmem, much more abundant than expected from the number of iBALT HA-specific GC B cells. This, together with our demonstration that lung Bmem exhibit a different transcriptional profile lead us to conclude that Bmem generated in other organs emigrate to lungs, as previously speculated by Allie *et al.* (2019). Here, we show that GC-derived Bmem in the lungs can be generated in spleen GC up to day 14 and mln GC at d28 and subsequently traffic to the lungs where they become tissue resident.

HA stem-specific Bmem have been described to derive from GC of lungs of IAV infected mice (Adachi et al., 2015). We did not assess stem *vs.* head specificity, but it is possible that iBALT derived Bmem harbor more cross-reactive B cells, particularly since their clonotypes do not overlap with mln-GC. Pulmonary Bmem also differed from spleen and mln in their transcriptional profiles, with upregulated *Cd69, Cd44, Ahr* and downregulated CD62L (*Sell*), *Cr2, Cd22*, among others. This confirms recent observations of organ-specific diversity of Bmem in bone marrow vs spleen (Riedel et al., 2020). It also highlights the complexity of the Bmem compartment, reflecting a need for specialization and rapid response of Bmem, depending on organ.

It is interesting to ponder the potential function(s) of early arriving Bmem cells in the lung. They may serve a back-up function to PB to insure clearance of the initital infection. They may also reflect early deployment against the chance of reinfection, particularly if niches for tissue resident Bmem are specifically created during the remodeling of the lung as the tissue returns to a pre-infection state. It is also possible that their function is antibody independent, e.g. participating in cytokine based tissue remodeling or immune cell regulation.

We observed a time dependent increase in BCR mutations in all B cell populations in all organs. Surprisingly, we did not detect a similar rate of clonal expansion in GC from mln and spleen, despite similar diversity and even after subsampling to equalize cell numbers. Clonal bursts (Tas et al., 2016) may be more common in mln because the total number of GC is lower or because of increased/persistent antigen levels. Alternatively, clonal expansion could be similar, but splenic GC may experience increased apoptosis. Whatever the explanation, the net result is the presence of a few hyperexpanded clonal families in mln GCs and many small clonal families in spleen GC.

Notably, previous studies using NP and HEL immunization model systems suggested a switch in the output of PB and Bmem, with early Bmem being unswitched, followed by swIg Bmem (between weeks 1-2) and then at day 21 the generation of PB (Weisel and Shlomchik, 2017). Except for early IgM Bmem, the response to IAV infection differs, featuring a constant output of PB and Bmem from GC, as judged by mutation rate. The strong correlation between clonal size and, importantly, mutational pattern suggests that Bmem are outputted constantly from GC.

Similarly, the origin of Bmem from GC has been hotly debated. Both inductive and stochastic models for Bmem differentiation have been proposed. Recently it has been proposed that Bmem precursors in the GC are selected into the memory compartment because of lower affintiy as compared to PB (Shinnakasu et al., 2016; Suan et al., 2017a). This notion is based on NP and HEL immunization using BCR transgenic mice. In both cases just one amino acid substitution is needed to dramatically improve BCR avidity (W33L for NP and Y53D for HEL). To relate this to anti-viral B cell responses we first examined the prototypical C12 idiotype in the anti-HA response, first described by Gerhard and colleagues, which is specific for the Cb antigenic site of HA (Kavaler et al., 1990). These cells carry a BCR with high germline affinity for HA, rapidly differentiate into extrafollicular plasma cells and do not participate in secondary responses to flu and therefore were assumed not to be forming memory (Kavaler et al., 1991; Rothaeusler and Baumgarth, 2010). Our analysis shows that these cells, of high affinity, are also capable of forming Bmem both as GC dependent as well as GC-independent.

Based on pseudotime analysis and comparison with previously identified gene signatures we identified PreMem-C9 as Bmem-precursor cells. Importantly, we did not find any significant differences in BCR that cells in this cluster expressed, neither in regard of mutational load, class switching, or clonality as compared to other GC cluster. 28 dpi GC-derived Bmem were at least as mutated as PB. While somatic hypermutation is not a perfect proxy for affinity it is suggestive that these cells underwent through similar number of selection cycles in the GC. Further, we detected a high proportion of cells expressing identical BCR in both Bmem and PB compartment which is again not compatible with affinity-based selection. This was further confirmed as we expressed mAbs from selected Bmem and PB and found their affinity to be comparable. Further, our results indicate the major determinant for affinity differences to be the clonal family. While a large number of Bmem that we detect are unmutated IgM, overall, the data presented here does not support the hypothesis of avidity-based selection for GC-derived Bmem, in case of an infection with a respiratory virus. It is possible that affinity selection is prevalent in immunization settings, where antigen amount is limiting but not when antigen is in excess, such as in the case of live replicating virus. In addition, soluble mAb expression might not fully recapitulate the complex GC environment, where avidity is also determined by multivalency and BCR density on B cells (Lingwood et al., 2012; Slifka and Amanna, 2019; Tolar and Pierce, 2010). Further, HA used for avidity measurements might not be in the same form presented in GC on FDC. Nevertheless, BCR from GC-derived Bmem and PB and even PreMem-C9 were indistinguishable by all the measures presented here. Our data suggests that stochasticity (Blink et al., 2005; Good-Jacobson and Shlomchik, 2010; Pelissier et al., 2020; Smith et al., 2000), might be one of the major determinants of Bmem differentiation from GC. Indeed, we identify clonal family as the main determinant of B cell affinity, a finding that would have been obviously impossible when using transgenic monoclonal mice. Importantly, we found the diversity of Bmem to be much larger than PB, with the latter mainly selected from, and composed of, expanded clones.

In summary, our study provides a comprehensive resource, linking BCR characteristics with transcriptional regulation of antigen specific B cell activation and differentiation, in different organs, upon respiratory virus infection.

## ACKNOWLEDGMENTS

We would like to thank the staff at the Experimental Biomedicine (EBM) core facility at the University of Gothenburg for animal management; E. Rekabdar and A. Almstedt, Genomics core facility at Sahlgrenska Hospital for running the sequencing and data pre-processing; M. Bäckstrom and R. Lymer, Mammalian Protein Expression (MPE) core facility at the University of Gothenburg for recombinant HA production and purification; D. Anastasakis, NIAMS, NIH, for help retrieving data sets for GSEA; A. Svitorka-Härtlova for assistance with figure creation with Biorender. D.A. is supported by the European Research Council starting grant (B-DOMINANCE, grant nr. 850638), the Swedish Research Council starting grant (grant nr. 2017-01439), the Jeanssons foundation (grant nr. JS2018-0011 and JS2019-0038), the Claes Groschinsky foundation (grant nr. M18237), the Institute of Biomedicine at the University of Gothenburg and the Bioinformatic long term support by the National Bioinformatics Infrastructure Sweden (NBIS) at SciLifeLab by Knut and Alice Wallenberg Foundation (grant nr. 1910). A.M.H. is supported by funding from the European Union’s Horizon 2020 research and innovation programme under SHIGETECVAX project (grant agreement No 815568) and VASA project (grant agreement No 815643), and the Innovative Medicines Initiative 2 Joint Undertaking under VSV-EBOPLUS project (grant agreement No 116068). V.G. is supported by the UiO World-Leading Research Community, the UiO:LifeSciences Convergence Environment Immunolingo and a EU Horizon 2020 iReceptorplus (#825821) project.

## AUTHOR CONTRIBUTIONS

NRM methodology, formal analysis, investigation, project administration, writing – review & editiing. JKJ data curation, methodology, formal analysis, software, writing – review & editing. IS methodology, investigation, writing – review & editing. JLR data curation, methodology, formal analysis, software, visualization, writing – review & editing. HA methodology, investigation, formal analysis. SSN methodology, investigation, formal analysis, writing – review & editing. AE methodology, investigation, writing – review & editing. PC data curation, methodology, formal analysis, software, writing – review & editing. WTY resources, writing – review & editing. CLF investigation. VB investigation. AMH resources, supervision, writing – review & editing. NL resources, supervision. NB resources, methodology, software, writing – review & editing. JWY resources, writing – review & editing. VG resources, methodology, software, writing – review & editing. MB methodology, formal analysis, investigation, writing – review & editing. DA conceptualization, data curation, formal analysis, funding acquisition, methodology, project administration, resources, software, supervision, visualization, writing – original draft, writing – review & editing.

## METHODS

### Mice and infection

C57BL/6 mice were purchased from Taconic Biosciences, Denmark and were housed in the animal facility of Experimental Biomedicine Unit at the University of Gothenburg. Female mice which are eight to twelve weeks old were used in the experiments. All the experiments were conducted according to the protocols (Ethical permit number: 1666/19) approved by regional animal ethics committee in Gothenburg. Mice were anesthetized with isoflurane and infected through nasal inoculation with 50 TCID_50_ Influenza A/Puerto Rico/8/34 (PR8) (Molecular clone; H1N1) diluted in HBBS containing 0.1% BSA.

### Cell sorting of Hemagglutinin-specific B cells

C57BL/6 mice were infected with PR8 H1N1 virus and were euthanized on different days post-infection. Lungs, spleen and mediastinal lymph nodes (mln) were isolated. The same organs from naïve mice were used as controls. Spleen and mln were mashed and passed through a 70µm filter to obtain single cell suspension. Lungs wereperfused and processed into single cell suspension using the mouse lung dissociation kit (Miltenyi Biotec) according to manufacturer’s instruction. Splenocytes and lung cells were enriched for total B cells using the EasySep Mouse Pan-B Cell Isolation kit (Stemcell Technologies) while whole mln cells were used for downstream processing. The cells were incubated for one hour at 4°C with a cocktail of fluorochrome-labelled antibodies consisting of anti-CD3-BV510 (cat. no: 563024, BD Biosciences), anti-B220-APC-Cy7 (cat. no: 552094, BD Biosciences) and anti-IgD-Pacific Blue (cat. no: 405712, BD Biosciences), and 1µg/ml biotinylated recombinant hemagglutinin (rHA) (Whittle et al., 2014) conjugated to streptavidin APC (cat. no: S868, Invitrogen). To exclude dead cells, the cells were washed and stained with LIVE/DEAD™ Fixable Aqua Dead Cell Stain (cat. no: L34957, Invitrogen) according to manufacturer’s instruction. A maximum of 10,000 live HA-specific mature B cells (CD3^-^B220^+^IgD^-^ rHA^+^) were sorted and collected in a BD FACSAria fusion or BD FACSAria III (BD Biosciences) cell sorter and processed immediately.

### Flow cytometry

All the fluorochrome-labelled antibodies used in flow cytometry were titrated for determining the optimal concentration. Briefly, spleens and lungs were harvested from C57BL/6 mice on day 14 post-PR8 H1N1 infection after euthanization. Spleens were processed into single cell suspension by mashing them and passing through a 70µm filter. Lungs were processed into single cell suspension using the mouse lung dissociation kit (Miltenyi Biotec). The following fluorochrome-conjugated antibodies were used for labelling the cells: anti-CD3-BV510 (cat. no: 563024, BD Biosciences), anti-B220-APC-Cy7 (cat. no: 552094, BD Biosciences), anti-NK1.1-BV510 (cat. no: 108737, Biolegend), anti-CD38-FITC (cat.no: 558813, BD Biosciences), anti-GL-7-PE-Cy7 (cat.no:144620,Biolegend),anti-IgD-BV786 (cat.no: 563618, BD Biosciences), anti-IgM-BUV395 (cat. no: 564025, BD Biosciences), anti-CD69-BUV737 (cat. no: 564684, BD Biosciences), anti-CD62L-BV711 (cat.no: 104445, Biolegend), anti-CCR7-BV605 (cat. no: 120125, Biolegend), anti-CD44- PE/Dazzle 594 (cat. no: 103055, Biolegend), anti-CD180-BV711 (cat. no: 740765, BD Biosciences), anti-CD22.2-BUV737 (cat. no: 741732, BD Biosciences), anti-CXCR4- PE/Dazzle 594 (cat. no: 146513, Biolegend), anti-CD83-PE (cat.no: 121507, Biolegend) and anti-CR2/CR1-BV421 (cat. no: 123421, Biolegend). The cells were stained with fluorochrome labelled antibodies for 20 min at 4°C. After washing, the cells are stained with LIVE/DEAD™ Fixable Aqua Dead Cell Stain (cat. no: L34957, Invitrogen) to exclude dead cells. The labelled cells were run and the data was acquired on the BD LSR Fortessa X-20 (BD Biosciences) and was analyzed using Flow Jo software (Tree Star).

### Generation and sequencing of single cell gene expression and enriched B cell libraries

Nearly 1500-10,000 sorted HA-specific mature B cells from individual organs were processed into single cells in a chromium controller (10X genomics). During this process, individual cells are embedded in Gel Beads-in-emulsion (GEMs) where all generated cDNA share a common 10X oligonucleotide barcode. After amplification of the cDNA, 5’gene expression library and enriched B cell library, with paired heavy and light chain were generated from cDNA of the same cell using Chromium single cell VDJ reagent kit (V1.1 chemistry, 10X genomics). The 5’gene expression libraries were sequenced in NextSeq or NovaSeq6000 sequencer (Illumina) using NextSeq 500/550 v2.5 sequencing reagent kit (read length: 2 × 75 bp) or NovaSeq S1 sequencing reagent kit (read length: 2 × 100 bp) (Illumina) respectively. The enriched B cell libraries were sequenced in NextSeq or MiSeq sequencer using NextSeq Mid Output v2.5 sequencing reagent kit (read length: 2 × 150 bp) or MiSeq Reagent Kit v2 (read length: 2 × 150 bp) (Illumina) respectively.

Lungs from mice M0_1, M7_2 and M14_1 and spleen from M28_3 failed to yield good quality GEMs and libraries and were not sequenced.

### Single-cell RNA-seq data processing

Single-cell RNA-seq data was processed in R with Sauron (https://github.com/NBISweden/sauron), which primarily utilizes the Seurat (v3.0.1) package (Stuart et al., 2019). This workflow comprises a generalized set of tools and commands to analyze single cell data in a more reproducible and standardized manner, either locally or in a computer cluster. The complete workflow and associated scripts are available on https://github.com/angelettilab/scMouseBcellFlu. A set of instructions on how to use the workflow and completely reproduce the results shown herein are available there.

Raw UMI count matrices generated from the cellranger 10X pipeline were loaded and merged into a single Seurat object. Cells were discarded if they met any one of the following criteria: percentage of mitochondrial counts > 25%; percentage of ribosomal (Rps or Rpl) counts > 25%; number of unique features or total counts was in the bottom or top 0.5% of all cells; number of unique features < 200; Gini or Simpson diversity index < 0.8. Furthermore, mitochondrial genes, non-protein-coding genes, and genes expressed in fewer than 5 cells were discarded, whereas the immunoglobulin genes *Ighd, Ighm, Ighg1, Ighg2c, Ighg2b, Ighg3, Igha*, and *Ighe* were retained in the dataset regardless of their properties.

Gene counts were normalized to the same total counts per cell (1000) and natural log transformed (after the addition of a pseudocount of 1). The normalized counts in each cell were mean-centered and scaled by their standard deviation, and the following variables were regressed out: number of features, percentage of mitochondrial counts, and the difference between the G2M and S phase scores.

Data integration across cells originating from different samples, time points and tissues were done on regressed scaled counts using the mutual nearest neighbors (MNN) (Haghverdi et al., 2018) on a set of highly variable genes (HVGs) identified within each sample individually and combined. The top 20 nearest neighbors (k) with a final dimensionality of 51 were used. Uniform Manifold Approximation and Projection (UMAP)(McInnes et al., 2018) was applied to the MNN-integrated data to further reduce dimensionality for visualization (2 dimensions) or for unsupervised clustering (10 dimensions).

At this stage, differential expression between clusters, and cell correlation with cell-type specific gene lists were evaluated to identify clusters of non-B-cells (such as NK or T cells). Predicted non-B-cells were removed from the data, and the entire single-cell RNA-seq processing pipeline was re-run using only the remaining B cells. Finally, hierarchical clustering was performed on the 10-dimensional UMAP embedding to define 16 clusters of B cell subtypes, which were then visualized on the 2-dimensional UMAP embedding.

### Trajectory inference analysis and RNA velocity

Trajectory inference analysis was performed on a diffusion map embedding (20 diffusion components; DCs) of the MNN-integrated count data using the destiny package (Angerer et al., 2016). The cell differentiation lineages were then predicted from the DCs using the slingshot package (Street et al., 2018). Cluster 3 (cells entering the GC) was specified as the starting point and cluster 5 as the end point (Bmem). Distance along the resulting curve was used to define the position of each cell in pseudotime. Identification of differentially expressed genes was done by fitting a generalized additive model (GAM) to the trajectory curve using the tradeSeq package (Van den Berge et al., 2019), allowing us to detect which genes exhibited expression behavior that was most strongly associated with progression along the defined lineage. Single-cell RNA-seq BAM files were processed using the velocyto command line tool (La Manno et al., 2018) to quantify the amount of unspliced and spliced RNA reads of each gene in each sample.

The scVelo package (Bergen et al., 2019) was used to perform the RNA velocity analysis. The first- and second-order moments for velocity estimation were calculated using the MNN-integrated data as the representation, and the cell velocities were computed using the likelihood-based dynamical model. A velocity graph was calculated based on cosine similarities between cells, and cell velocities were visualized as streamlines overlaid on the 2-dimensional UMAP embedding.

### BCR sequence data processing

The BCR sequence data was processed using the Immcantation toolbox (v4.0.0) using the IgBLAST and IMGT germline sequence databases, with default parameter values unless otherwise noted. The IgBLAST database was used to assign V(D)J gene annotations to the BCR FASTA files for each sample using the Change-O package (Gupta et al., 2015), resulting in a matrix containing sequence alignment information for each sample for both light and heavy chain sequences.

BCR sequence database files associated with the same individual (mouse) were combined and processed to infer the genotype using the TIgGER package (Gadala-Maria et al., 2015) as well as to correct allele calls based on the inferred genotype. The SHazaM package (Gupta et al., 2015) was used to evaluate sequence similarities based on their Hamming distance and estimate the distance threshold separating clonally related from unrelated sequences. The predicted thresholds ranged from 0.096 to 0.169, where a default value of 0.1 was assumed for cases when the automatic threshold detection failed. Ig sequences were assigned to clones using Change-O, where the distance threshold was set to the corresponding value predicted with SHazaM in the previous step. Germline sequences were generated for each mouse using the genotyped sequences (FASTA files) obtained using TIgGER (Gadala-Maria et al., 2015). BCR mutation frequencies were then estimated using SHazaM. The BCR sequence data, clone assignments, and estimated mutation frequencies were integrated with the single-cell RNA-seq data by aligning and merging the data with the metadata slot in the processed RNA-seq Seurat object.

### Identification of clones and diversity using scRepertoire

scRepertoire (Borcherding et al., 2020) was used to determine clonal groups based on paired heavy and light chains. This package uses the filtered contig annotation obtained from cell ranger. Clones were assigned only for cells were high quality paired heavy and light chains were sequenced. Clones were assigned based on the CTstrict function per each mouse. The CTstrict function consider clonally related two sequences with identical V gene usage and >85% normalized hamming distance of the nucleotide sequence. Percent of unique clonotypes were obtained using the quantContig function. Integration with the Seurat object was done using the combineExpression function. Ranking of clones were determined using the clonalProportion function and Shannon and Simpson’s diversity determined using the clonalDiversity function. All function were run using the exportTable = T function to obtain a table and customarily facet the graph in R using the ggplot package. Sharing of clones between clusters was visualized using the ggalluvial package.

### Differential gene expression analysis

Differentially expressed genes between different clusters, organs, isotype or differentially mutated cells were identified using the FindAllMarkers function from Seurat using default settings (Wilcoxon test and Bonferroni *p* value correction). Significant genes with average log fold change > 0.25 and expressed in >25% of cells in that group were ranked according to fold change and represented in the FeaturePlot.

### Gene set enrichment analysis (GSEA)

For GSEA analysis, differentially expressed genes for each cluster or organ were calculated using the Wilcoxon rank sum test via the wilcoxauc function of the presto package using default parameters (including Benjamini-Hochberg false discovery rate correction) and filtered on logFC >1 and *padj* <0.05. GSEA was run on pre-ranked genes using the fgsea package (Korotkevich et al., 2019).For each enrichment graph we report *p, padj* (FDR *q*) and NES (enrichment score normalized to mean enrichment of random samples of the same size) values in the figure.

### Generation of clonal trees and expression of monoclonal antibodies

Five clonal families were randomly selected among the hyperexpanded, as defined by scRepertoire. Clonal trees were reconstructed using the Alakazam package of Immcantation (Gupta et al., 2015). In brief, clones were made with the function makeChangeOclones and lineages were reconstructed using the dnapars function of the Phylip package via buildPhylipLineage function. Clonal trees were visualized via the iGraph package in R. Random representative clones were selected from each family. Both heavy and light chain sequences were synthesized (Twist Bioscience) and subsequently cloned into a mouse IgG1 expression vector (Kosik et al., 2019). To confirm the cloning developed vectors were sequenced by Eurofins Genomics. To express recombinant antibodies plasmids encoding corresponding heavy and light chains were mixed in equal ratio. Transfection of Expi293 cells was carried out by ExpiFectamine 293 Transfection Kit (ThermoFisher) according to the manufacturer instruction. After four days supernatants were collected and filtered. Purification of the immunoglobulins was carried out by Akta Start System (GE Healthcare) using protein G column. Elution of bound antibodies was done by 0.1 M glycine buffer, pH 2.7. To neutralize the solution coming from the column collecting tubes contained 1 M Tris buffer, pH 9.0. Antibody-containing eluates were concentrated by centrifugation through VivaSpin columns with a 30 kDa cut-off. Estimation of antibody concentration was done by NanoDrop (ThemoFisher) equipment. Binding of all mAbs was confirmed using ELISA. Briefly, plates were coated overnight with 20HAU of PR8 virus in PBS. Plates were blocked with PBS 2% milk for 1h at RT. mAbs were serially diluted and incubated for 1hr at RT. After washing, plates were incubated with anti-mouse IgG secondary Ab (Southern Biotech, cat no 1170-05, dilution 1:1000) 1hr at RT. Plates were developed with 1-step Ultra TMB-ELISA (ThermoFisher), reaction stopped by 2M H_2_SO_4_ and read at 450nm.

### Bio-layer interferometry

Biolayer Interferometry (BLI)-based assay was set up to measure the affinity of murine mAbs to Pro-haemeagglutinin (HA). In principle, BLI is an optical analytical, label-free, technique and is used to analyze biological interactions using the difference in interference pattern of white light reflected from two surfaces: a layer of immobilize ligand on the biosensor tip and an internal reference (blank) layer.

Kinetic assays were optimized for buffer, pH, temperature conditions, orbital shake speed for affinity analysis. Briefly, 0.5 µg of AviTagged HA diluted in acetate 4.5 (pall fortebio, Sartorius group, Amsterdam, Netherlands) was immobilized onto SAX biosensors (high precision streptavidin sensors: Pall fortebio, Sartorius group, Amsterdam, Netherlands). mAbs were diluted 1:1000 ratio in kinetic buffer (1%BSA in PBS, pH 7.5) Pall fortebio, Sartorius group, Amsterdam, Netherlands). H26A1, H28-E23 and H17-40 mAbs were used as positive controls.

Kinetics of binding interactions of mAbs to the immobilized HA were determined using octet data acquisition software (version 10.0.087) blank experiment with the following experimental steps: wash (PBS buffer, pH 7.5; 60 seconds), immobilization (HA, 0.5 µg; 300 seconds), association (analytes, 1:1000 of mAbs; 420 seconds), dissociation (PBS buffer, pH 7.5; 600 seconds), regeneration (10mM glycine, sigma-aldrich, Stockholm, Sweden; 60 seconds), wash (PBS buffer, pH 7.5; 60 seconds). Experiments were carried out at plate shake speed of 1000 rpm and plate temperature of 25 °C. Reference sensor was immobilized with PBS (pH 7.5, Gibco, Sweden) and samples were run with same experimental conditions as ligand immobilized sensor. Data was processed using octet data analysis software (version 10.0) and on-rate (ka), off-rate (kd) and affinity (KD) were calculated upon reference subtraction.

### Statistical analysis

For single cells analyses statistics are described above. For multiple comparison between groups, One-way ANOVA or two-way ANOVA with Tukey’s multiple comparisons were used.

### Code availability

The complete workflow and associated scripts are available on https://github.com/angelettilab/scMouseBcellFlu. A set of instructions on how to use the workflow and completely reproduce the results shown herein are available there.

